# Formation and disassembly of a contractile actomyosin network mediates content release from large secretory vesicles

**DOI:** 10.1101/216044

**Authors:** Dagan Segal, Assaf Zaritsky, Eyal D. Schejter, Ben-Zion Shilo

## Abstract

Secretion of adhesive glycoproteins to the lumen of *Drosophila* larval salivary glands is carried out by contraction of an actomyosin network that is assembled around large secretory vesicles, following their fusion to the apical membranes. We have identified a cycle of actin coat nucleation and disassembly that is independent of myosin. Recruitment of active Rho1 to the fused vesicle triggers activation of the formin Diaphanous and nucleation of linear actin. This, in turn, leads to actin-dependent localization of a RhoGAP protein that locally shuts off Rho1, promoting disassembly of the actin coat. Recruitment of the branched actin nucleation machinery is also required for effective Rho1 inactivation. Interestingly, different blocks to actin coat disassembly arrested vesicle contraction, indicating that actin turnover is an integral part of the actomyosin contraction cycle. The capacity of F-actin to trigger a negative feedback on its own production may be utilized in a variety of scenarios, to coordinate a succession of morphogenetic events or maintain homeostasis.

**Summary:** This work identified a cycle of actin assembly and disassembly in large secretory vesicles of *Drosophila* salivary glands. Actin disassembly is triggered by actin-dependent recruitment of a RhoGAP protein, and is essential for the contractility of the vesicle leading to content release to the lumen.

## Introduction

Coordinated formation and disassembly of contractile actin-based structures has been shown to underlie diverse settings of tissue morphogenesis. These events are not regulated through gene expression, but rather rely on inherent mechanistic features, as well as responses to tension and stress (Guillot and Lecuit, 2013; Pandya et al., 2017). The tight temporal and spatial regulation of actomyosin dynamics, and the versatile activity of the same basic machinery in diverse biological scenarios, remains to be fully understood.

Exocytosis from the epithelium of *Drosophila* larval salivary glands provides a powerful system to study the orchestrated regulation of actomyosin during contractile events (reviewed in (Tran and Ten Hagen, 2017)). Upon stimulation by hormonal cues, these cells secrete large amounts of “glue” glycoproteins from their apical membrane into the lumen of the gland over the course of two hours, in preparation for external deposition that will enable the larva to adhere to a dry surface before pupariation (Biyasheva et al., 2001). The glue proteins are stored prior to secretion in very large vesicles, ranging in size from 4 to 8 microns in diameter. Content release from these vesicles into the lumen is achieved following their fusion with the apical membrane of the cell, through formation of an actin coat around each vesicle and recruitment of myosin, which together mediate the forces necessary for vesicle contraction (Rousso et al., 2016; Tran et al., 2015) (Figure 1a-b).

**Figure 1:**
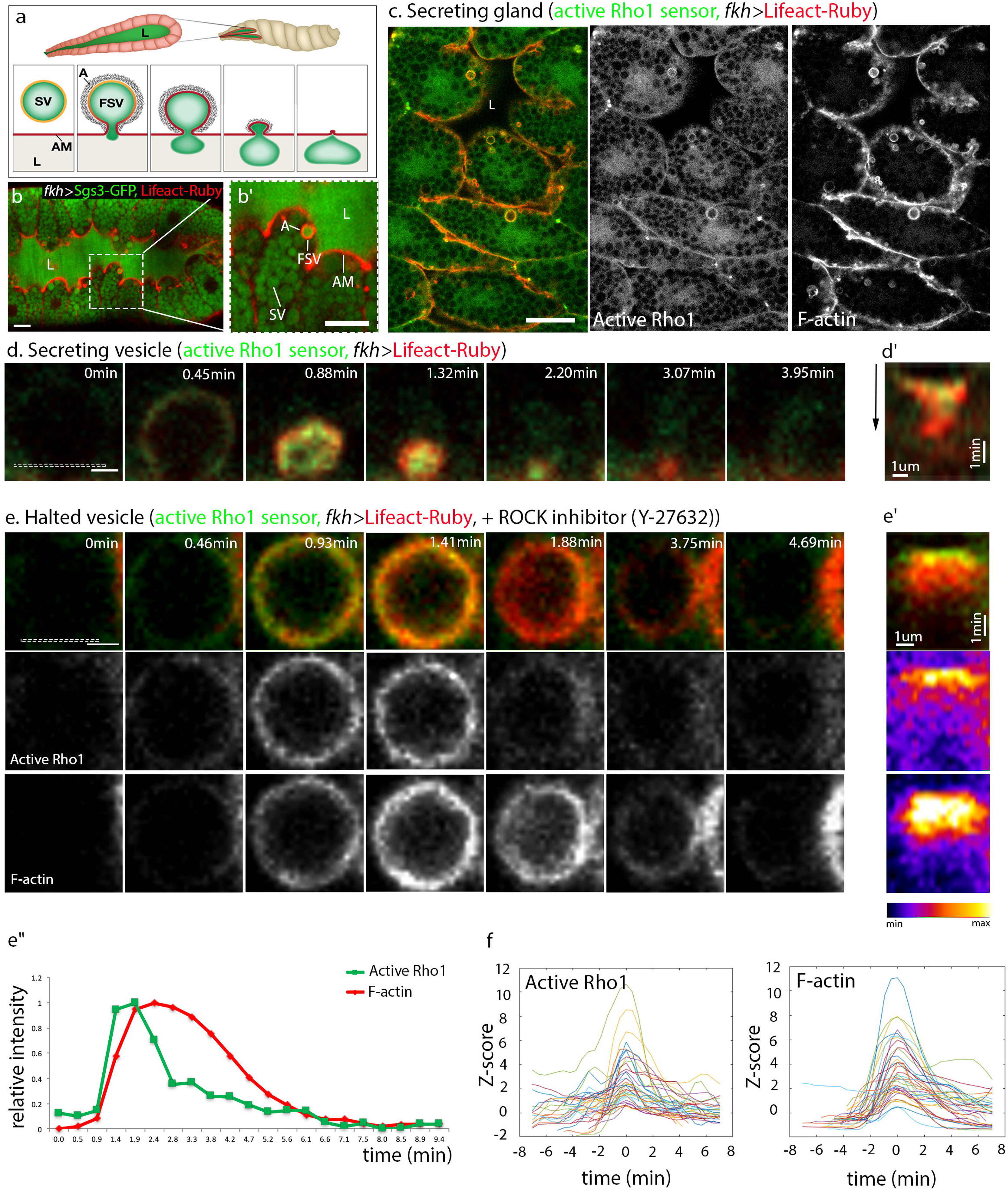
Actin recruitment and disassembly are driven by the activity of Rho1. **(a-b’)** Schematic drawing **(a)** and confocal microscope image **(b)** of a *Drosophila* third instar salivary gland expressing the glue protein Sgs3-GFP (green) and the F-actin marker Lifeact-Ruby (red, A). **(b’)** shows enlargement of boxed area in **(b)**. Large secretory cells surround an expanding lumen (L), and are filled with secretory vesicles (SV) containing the glue cargo. During secretion, vesicles fuse to the actin-rich apical membrane (AM) facing the lumen. Fused secretory vesicles (FSV) form an actomyosin coat (A) that enables contraction and expulsion of their content into the lumen**. (c)** A secreting gland displays active Rho1 (visualized by the sensor Ani-RBD-GFP(Munjal et al., 2015), green) and F-actin (red) enriched on vesicles that fused to the apical membrane, at various stages of contraction. **(d)** Time course of a single secreting vesicle. The vesicle contracts fully within three minutes, with residual actin cleared from the site of fusion after ~1 additional minute. **(d’)** Kymograph monitoring active Rho1 and F-actin dynamics at the vesicle/plasma membrane interface (dashed-box area in initial panel of **(d)**). Time progresses downwards as indicated by black arrow. Dynamics of actin clearing are difficult to characterize due to drastic morphological changes. **(e)** Addition of the ROCK inhibitor Y-27632 causes accumulation of contraction-halted vesicles, allowing unperturbed visualization of active Rho1 and actin dynamics (see also Movie S1). Halted vesicles exhibit active Rho1 and actin coat enrichment, followed by Rho1 inactivation and actin coat disassembly, in a timeframe comparable to that of a squeezing vesicle. **(e’)** Kymographs monitoring active Rho1 and F-actin dynamics at the vesicle surface (dashed-box area in initial panel of **(e)**)**. (e’’)** Graph plotting relative intensity of active Rho1 (green) and F-actin (red) in the area monitored in the kymographs **(e’)** over time. Intensity is normalized to the highest value within each channel. Note that both active Rho1 and actin rise sharply, but actin disassembly is gradual while Rho inactivation is sharp. **(f)** Dynamic profiles over time of active Rho1 and F-actin in 38 vesicles from a single contraction-halted gland. Vesicle timescales are aligned to intensity peaks. Z-score is defined as the number of standard deviations that the signal intensity differs from background levels. The duration and dynamics of active Rho1 and actin are highly similar between vesicles, despite large variations in Z-score. Scale bars: 20μm in b, b’, c. 2μm in d and e.

The basic molecular network underlying this actomyosin-dependent secretion cycle, which lasts 1-3 minutes, has been established (Rousso et al., 2016; Tran et al., 2015): Upon fusion with the apical membrane, activated Rho1-GTP is enriched on the fused vesicle, and subsequently recruits and activates the formin Diaphanous (Dia), that in turn generates a linear actin coat around the vesicle. In parallel, Rho1 triggers Rho kinase (ROCK; Rok in *Drosophila*), which recruits myosin unto the actin coat and mediates contraction. Incorporation of the branched actin nucleation machinery to the fused vesicle is also critical for actomyosin contractility. Each vesicle releases its content within approximately 4 minutes after fusion, and the dense actomyosin network surrounding it disintegrates. Thus, every secretory glue vesicle presents an entire cycle of actomyosin coat assembly and disassembly, providing a unique system to identify the relevant molecular players and to dissect their regulatory interactions.

Surprisingly, we have found that disassembly of the actin coat, which accompanies vesicle content release, is necessary for contraction of the actomyosin network. A dedicated Rho GTPase activating protein (RhoGAP) mediates Rho1 inactivation, leading to actin disassembly. RhoGAP activity and recruitment to the contracting vesicle appear to depend on F-actin, while branched-actin polymerization is necessary for RhoGAP activity but not its recruitment. The sequential temporal recruitment of active Rho1 and its inhibitors is made evident by the multiple cycles of accumulation and dispersion of active Rho1 and the actin coat observed on individual fused but contraction-halted vesicles, implying a negative-feedback-based mechanism. Actin coat dynamics and the resulting vesicle contraction are therefore achieved by coordinating formation and disassembly of the actomyosin network.

## Results

### Actin recruitment and disassembly is driven by the activity of Rho1

We examined the dynamics of actin recruitment and its regulators on secretory vesicles through live, *ex-vivo* imaging of the *Drosophila* salivary gland. Previously, we have shown that stimulation of Dia nucleation activity by active Rho1 is required for generation of the bulk of the actin coat surrounding the secretory vesicles (Rousso et al., 2016). Myosin II is also recruited to the vesicle in a Rho1/Rok-dependent manner (Rousso et al., 2016), thereby forming the actomyosin network necessary for vesicle contraction. Concomitant with squeezing of the vesicle, actin as well as active Rho1 are cleared from the site of fusion (Figure 1c, d).

The rapid reduction in size of the vesicle as it contracts, and the enriched levels of filamentous actin continuously present on the apical membrane, make it difficult to reproducibly follow the dynamics of actin disassembly on the vesicle. We therefore used a chemical inhibitor of ROCK kinase activity (Y-27632)(Uehata et al., 1997) to hinder Myosin II activation, with the aim of preventing and uncoupling vesicle contraction from actin coat disassembly, thereby enabling clearer visualization and quantification of actin dynamics on the vesicles that fail to squeeze. As expected, Myosin II is not recruited to vesicles upon addition of Y-27632 (Figure S1). While these vesicles still fuse with the apical membrane and an actin coat is formed around them, they do not contract (Figure 1e, movie S1). The contraction-halted vesicles display a clear disassembly of actin, which, importantly, exhibits a time course that is comparable to that of a squeezing vesicle (Figure 1d). Similar effects have been observed upon knockdown of the *Drosophila* Myosin II (heavy chain) homolog *zipper* (Rousso et al., 2016). These features make the halted vesicles a useful setting for studying actin coat dynamics, and, furthermore, demonstrate that actin disassembly is independent of Myosin II or vesicle contraction.

Monitoring of the Ani-RBD-GFP sensor revealed that active Rho1 levels associated with the halted vesicles not only rise but also diminish, and do so in concert with the dynamics of actin coat assembly and disassembly (Figure 1e). Semi-automated quantification on multiple vesicles within a single gland (Figure 1f) or multiple glands (Figure S2, see methods) allowed further characterization of these active Rho1 and actin cycles. Cycle duration commonly ranged between 1-3 minutes for active Rho1, and 3-7 minutes for actin (Figure S2a), despite large variations in signal amplitude (Figure 1f). This is comparable to the time scale of content release by a naturally contracting vesicle, which takes place over 1-3 minutes, with subsequent actin clearing taking roughly one additional minute (Rousso et al., 2016) (Figure 1d). More detailed analysis showed that Rho1 deactivation precedes actin coat disassembly (Figure S2b). These quantifications suggest that all features of the actin cycle are dependent on Rho1 activity: actin coat assembly is triggered (via Dia) following Rho1 activation (Rousso et al., 2016), while coat depletion follows shutdown of Rho1 activation and a resulting shift towards filament depolymerization.

### RhoGAP71E is required for Rho1 inactivation

Since Rho1 inactivation precedes and likely leads to actin coat disassembly, we examined the potential involvement of Rho GTPase activating proteins (RhoGAPs), which commonly facilitate the conversion of RhoGTPases from an active GTP-bound to an inactive GDP-bound state (Tcherkezian and Lamarche-Vane, 2007). While *Drosophila* has over 20 RhoGAP proteins (Greenberg and Hatini, 2011), RNA-seq data suggests that only a subset of eight RhoGAPs are expressed in the salivary gland at high or moderate levels (Brown et al., 2014). We conducted a functional knockdown screen, by monitoring active Rho1 sensor and actin coat dynamics on contraction-halted vesicles, following salivary-gland expression of RNAi constructs directed against each of the eight RhoGAP genes (table S1).

A unique pattern of sustained Rho1 activation and a persistent actin coat, the predicted phenotype for interference with Rho1 inactivation, was observed following knockdown of only one of these elements, RhoGAP71E (also termed C-GAP (Mason et al., 2016)). Y-27632-treated salivary glands expressing *RhoGAP71E* RNAi driven by *fkh-*GAL4 displayed vesicles on which both the active Rho1 sensor and the actin coat were sustained for several minutes (Figure 2a-b, Figure S2c). Three different RNA-i constructs targeted against RhoGAP71E showed similar results (Table S1). In contrast, the active Rho1/actin cycles on contraction-halted vesicles were not affected following expression of all other RhoGAP RNAi constructs (for example, Figure S3a).

**Figure 2:**
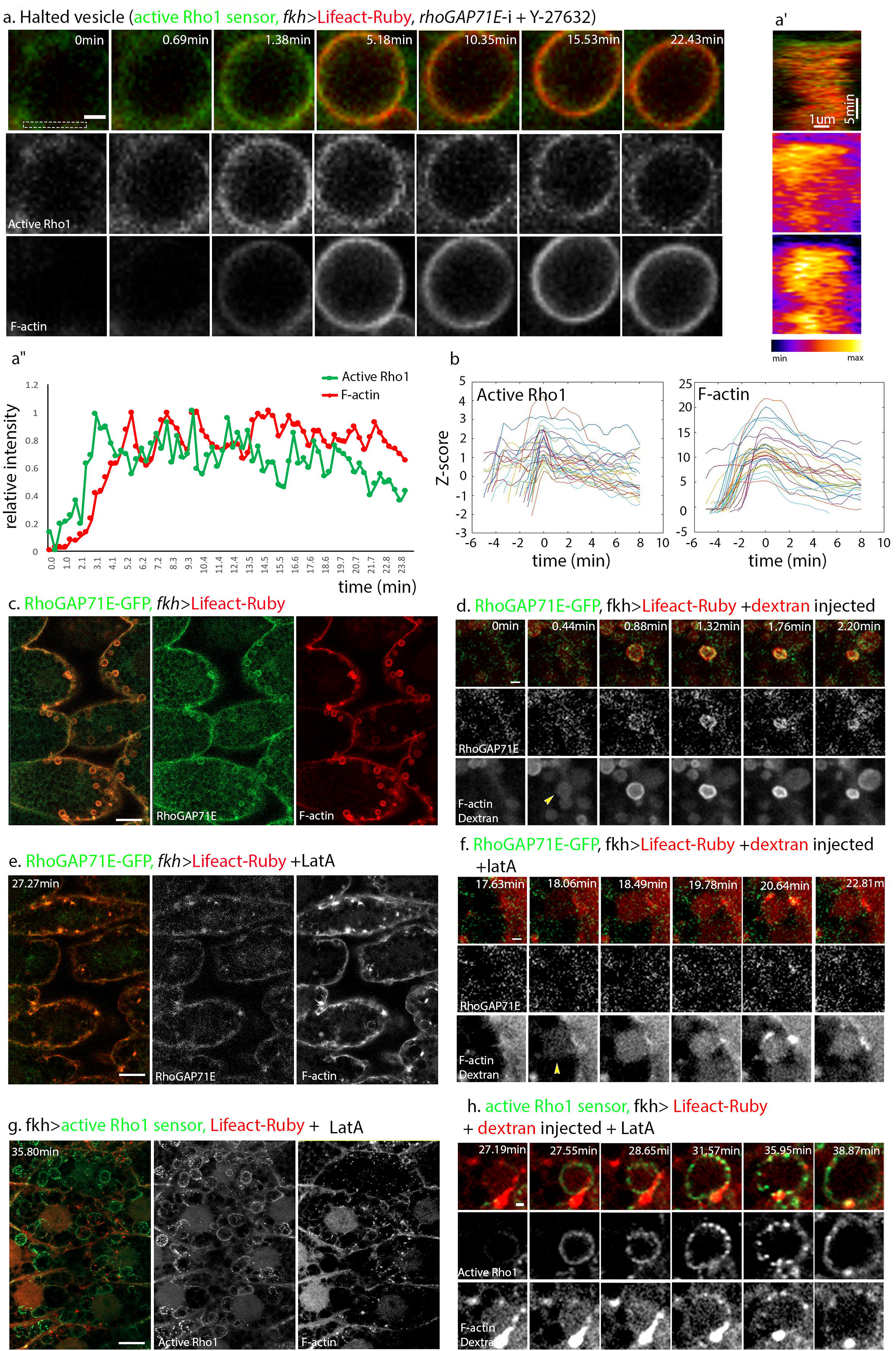
RhoGAP71E is required for Rho1 inactivation. A candidate based RNA-i screen identified RhoGAP71E as a regulator of Rho1 inactivation in the salivary gland. **(a)** Contraction-halted glands expressing the sensor for active Rho1 (green) and Lifeact-Ruby (red) along with *RhoGAP71E* RNAi show sustained active Rho1 and actin coat. **(a’)** Kymographs monitoring active Rho1 and F-actin dynamics at the vesicle surface (dashed-box area in initial panel of **(a)**). **(a’’)** Graph plotting relative intensity of active Rho1 (green) and F-actin (red) in the area monitored in the kymographs **(a’)** over time. Intensity is normalized to the highest value within each channel. Note that both active Rho1 and actin rise sharply and are sustained on the vesicle for over 20 minutes. **(b)** Dynamic profiles over time of active Rho1 and F-actin in 29 vesicles from a single contraction-halted gland. Vesicle timescales are aligned to intensity peaks. Z-score is defined as the number of standard deviations that the signal intensity differs from background levels. Note that while Rho1 and actin rise sharply, they are sustained above initial levels over many minutes (compare timescale to Figure 1f). **(c)** Glands expressing RhoGAP71E-GFP, an endogenously expressed GFP-tagged version of RhoGAP71E (green) and Lifeact-Ruby (red). RhoGAP71E-GFP appears to be enriched in most actin-coated vesicles. **(d)** A single secreting vesicle in such a gland expressing RhoGAP71E-GFP (green) and Lifeact-Ruby (red), with TMR-dextran (red) injected into the lumen to mark fused vesicles. Yellow arrowhead marks appearance of dextran within the vesicle, preceding formation of the bright actin coat on the vesicle surface. RhoGAP71E-GFP and actin appear to be simultaneously enriched on vesicles within 30 seconds after vesicle fusion. **(e)** Glands expressing RhoGAP71E-GFP (green) and Lifeact-Ruby (red). Addition of the F-actin polymerization inhibitor Latrunculin-A (LatA) to the medium caused loss of both actin coat formation and RhoGAP71E recruitment to vesicles. Time indicated corresponds to time elapsed since LatA addition. **(f)** A single secreting vesicle in such a LatA-treated gland expressing RhoGAP71E-GFP (green) and Lifeact-Ruby (red), injected with TMR-dextran (red) into the lumen. Yellow arrowhead marks appearance of dextran and onset of fusion. While vesicles still fuse after 17 minutes of LatA addition, actin and RhoGAP71E-GFP are not recruited to newly fused vesicles, demonstrating that RhoGAP71E recruitment requires F-actin. **(g)** LatA-treated glands expressing the active Rho1 sensor (green) and Lifeact-Ruby (red). Unlike RhoGAP71E-GFP, active Rho1 is enriched on a large subset of vesicles, despite the failed recruitment of actin. **(h)** A single secreting vesicle in such a Lat-A-treated gland expressing RhoGAP71E-GFP (green) and Lifeact-Ruby (red), and injected with TMR-dextran (red) into the lumen. Active Rho1 is recruited and sustained upon the fused vesicle, with a progressively punctate pattern forming over time. In addition, compound vesicle fusion leads to formation of abnormally large vesicles. Under these conditions, active Rho1 is sustained for at least 10 minutes in 42±19% of vesicles (n=67, measured in 3 of 5 glands which displayed this phenomenon), implying lack of Rho1 inactivation in the absence of actin and RhoGAP71E. Scale bars: 2μm in a, d, f, h. 20μm in c, e, g.

RhoGAP71E has been implicated in spatial restriction of Rho1 activity and subsequent cell contraction during gastrulation of *Drosophila* embryos (Mason et al., 2016). To follow the role of RhoGAP71E in regulation of Rho1 activity in glue vesicles, we sought to examine its dynamic localization on the secretory vesicles. Towards this end, we created an endogenously expressed GFP-tagged version of RhoGAP71E (RhoGAP71E-GFP), using the RMCE technique (Venken et al., 2011). RhoGAP71E-GFP is recruited to most fused vesicles displaying an actin coat (Figure 2c). We monitored the onset of vesicle fusion with the apical membrane by injection of fluorescent Dextran into the salivary gland lumen, since the dye enters the vesicle immediately upon fusion (Rousso et al., 2016). Using this method, we followed the temporal dynamics of RhoGAP71E-GFP during secretion, which appears to be recruited to the vesicles after fusion and concurrently with assembly of the actin coat (Figure 2d). Subsequent higher time-resolution imaging showed that RhoGAP is, in fact, recruited slightly after the actin coat (Figure S3c). RhoGAP71E is removed once the actin coat is disassembled, as observed in halted vesicles (Figure S3b).

Recent work in the early *C. elegans* embryo has suggested that actin can serve as a recruiting element for RhoGAPs (Robin et al., 2016). To determine whether actin plays a similar role in the context of glue vesicle secretion, we examined the effects of inhibition of actin polymerization, using the chemical inhibitor Latrunculin-A (LatA). Actin-coated vesicles are no longer observed within 20 minutes of LatA addition (Figure 2e). Dextran injection to the lumen of such LatA-treated glands demonstrates that vesicle fusion is unaffected (Figure 2f), although, as expected, these vesicles do not contract since they lack an actomyosin coat. Active Rho1 is recruited to these vesicles, albeit with a somewhat irregular pattern (Figure 2g, h). This suggests that enrichment of active Rho1 is independent of actin coat formation, consistent with its role in triggering actin polymerization. In contrast, RhoGAP71E-GFP recruitment was abolished entirely in newly fused vesicles lacking an actin coat (Figure 2e, f). Correspondingly, prolonged incubation with Lat-A (> 30 minutes) resulted in sustainment of active Rho1 for at least 10 min in 42±19% of vesicles (n=67, Figure 2h, Movie S2).

Taken together, these results suggest that while Rho1 activation is independent of F-actin, RhoGAP71E recruitment and subsequent Rho1 inactivation are F-actin dependent.

### Coordinated oscillations of active Rho1, actin, and RhoGAP71E

Strikingly, prolonged tracking of individual contraction-halted vesicles uncovered coordinated oscillations of active Rho1 and F-actin upon a large subset of halted vesicles (Figure 3a-a”). Since these vesicles continue to be fused to the apical membrane, which provides a permanent source of active Rho1 (Rousso et al., 2013), the observed additional cycles of actin coating upon the same vesicle may be a consequence the of initial disassembly of the actin coat and the concomitant loss of RhoGAP71E (see Figure 2). Automated scoring of vesicles for oscillations (see supplemental methods) showed that 52.9% of vesicles that were tracked for 15.5 minutes or more displayed at least one oscillation in addition to the initial complete actin cycle, with several observable cases of two or more oscillations upon the same vesicle (for example, Figure S4a). The median time interval between oscillations was approximately 6 minutes (Figure 3b), which is similar in timescale to a single actin cycle (Figure S2). Notably, instances of oscillations of actin can also be observed upon contracting vesicles, although these events require an increased temporal resolution and are challenging to resolve from the cortical actin associated with the apical membrane (Figure S4b).

**Figure 3:**
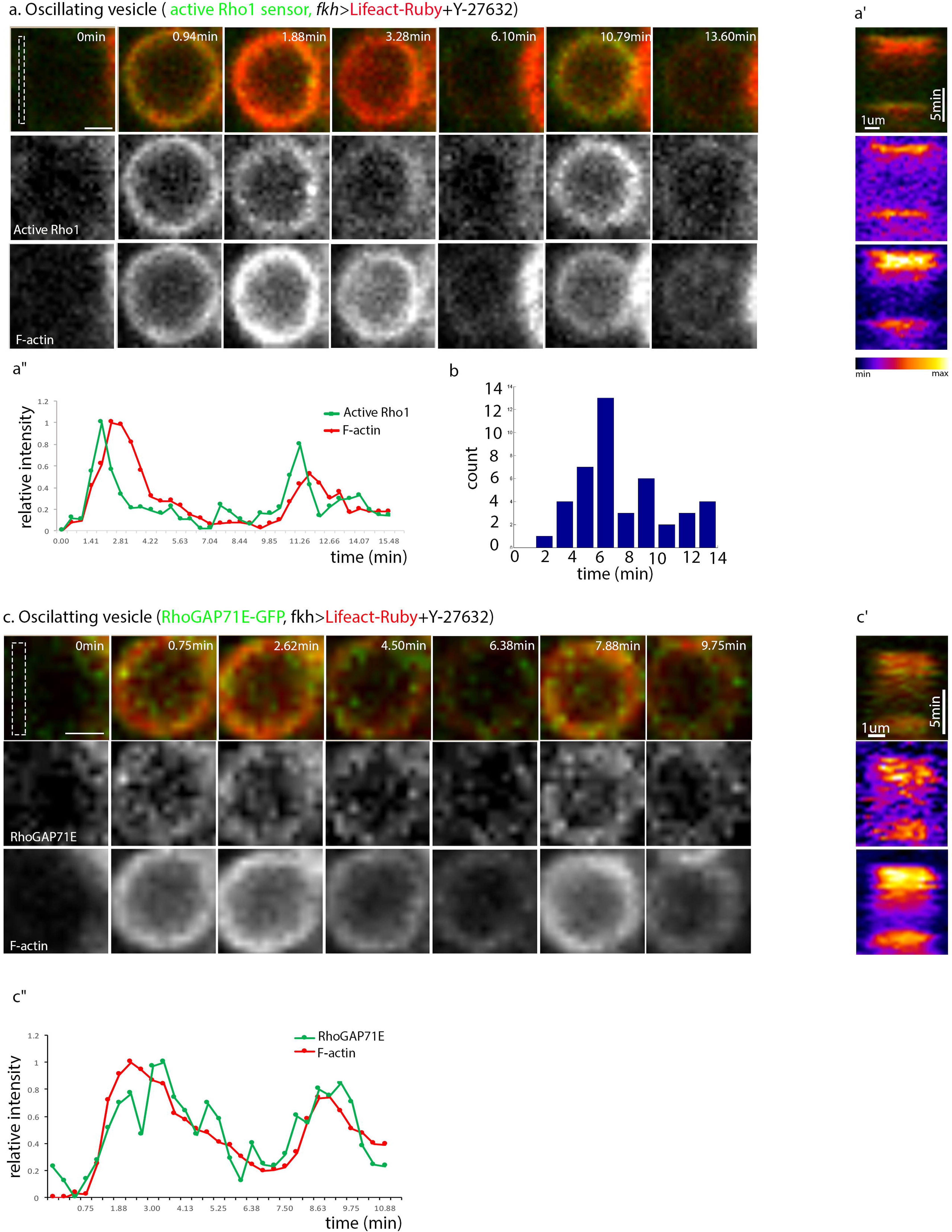
Coordinated oscillations of active Rho1, F-actin, and RhoGAP71E. **(a)** Coordinated oscillatory behavior of the active Rho1 sensor (green) and Lifeact-Ruby (red) on a single contraction-halted vesicle from a Y-27632-treated salivary gland. **(a’)** Kymograph monitoring active Rho1 and F-actin dynamics at the vesicle surface (dashed-box area in initial panel of **(a)**) **(a’’)** Graph plotting relative intensity of active Rho1 (green) and F-actin (red) in the area monitored in the kymograph **(a’)** over time. Intensity is normalized to the highest value within each channel. **(b)** Histogram showing the time in minutes between peaks of actin accumulation in oscillating vesicles (n=43 in 4 glands). Oscillations most frequently reach their maximum ~6 minutes after the previous peak, similar to the time scale of a single actin cycle. Out of vesicles that were tracked for at least 15.5 minutes (defined by 90^th^ percentile of oscillation peaks +3 seconds), 52.9% displayed at least one oscillation in addition to their first peak (n= 34 in 4 glands). **(c)** Coordinated oscillatory behavior of RhoGAP71E-GFP (green) and Lifeact-Ruby (red) on a single contraction-halted vesicle from a Y-27632-treated salivary gland. **(c’)** Kymograph monitoring RhoGAP71E-GFP and F-actin dynamics at the vesicle surface (dashed-box area in initial panel of **(c)**). **(c’’)** Graph plotting relative intensity of RhoGAP71E-GFP (green) and F-actin (red) in the area monitored in the kymograph **(c’)** over time. Intensity is normalized to the highest value within each channel. Note that RhoGAP71E-GFP and actin appear and disappear in a similar temporal pattern, consistent with a role for F-actin in RhoGAP71E recruitment. Scale bars: 2μm in a, c.

Importantly, RhoGAP71E-GFP recruitment to contraction-halted vesicles also exhibits an oscillatory behavior, in apparent coordination with the repeated actin cycles (Figure 3c). Taken together, these results suggest that Rho1, the actin coat, and RhoGAP71E are all part of a shared regulatory circuitry, and are sequentially recruited. This temporally resolved regulatory cycle appears to be initiated by Rho1 activation, which subsequently recruits actin and RhoGAP71E to the vesicle, and ultimately leads to Rho1 inactivation and actin disassembly with a temporal delay comparable to the time of a squeezing vesicle.

### Actin disassembly is essential for vesicle contraction

We next sought to assess the role of the Rho1-actin-RhoGAP71E regulatory network, which we had identified on contraction-halted vesicles, in the context of normal glue content release. While formation of the Rho1-dependent actomyosin coat is essential for vesicle contraction, the significance of coat disassembly is not immediately apparent. We first addressed this issue via knockdown of *RhoGAP71E*. Strikingly, this treatment resulted-on its own-in an arrest of secretory vesicle contraction, as seen by accumulation of halted vesicles in the entire gland (Figure 4a, Movie S3), and failed vesicle constriction (Figure 4b). Both active Rho1 and the actin coat persist on these vesicles, similarly to the effect of *RhoGAP71E* knockdown on contraction-halted (Y-27632-treated) vesicles (see Figure 2). These observations suggest, therefore, that Rho1 inactivation and the subsequent actin turnover are essential for proper actomyosin contraction.

**Figure 4:**
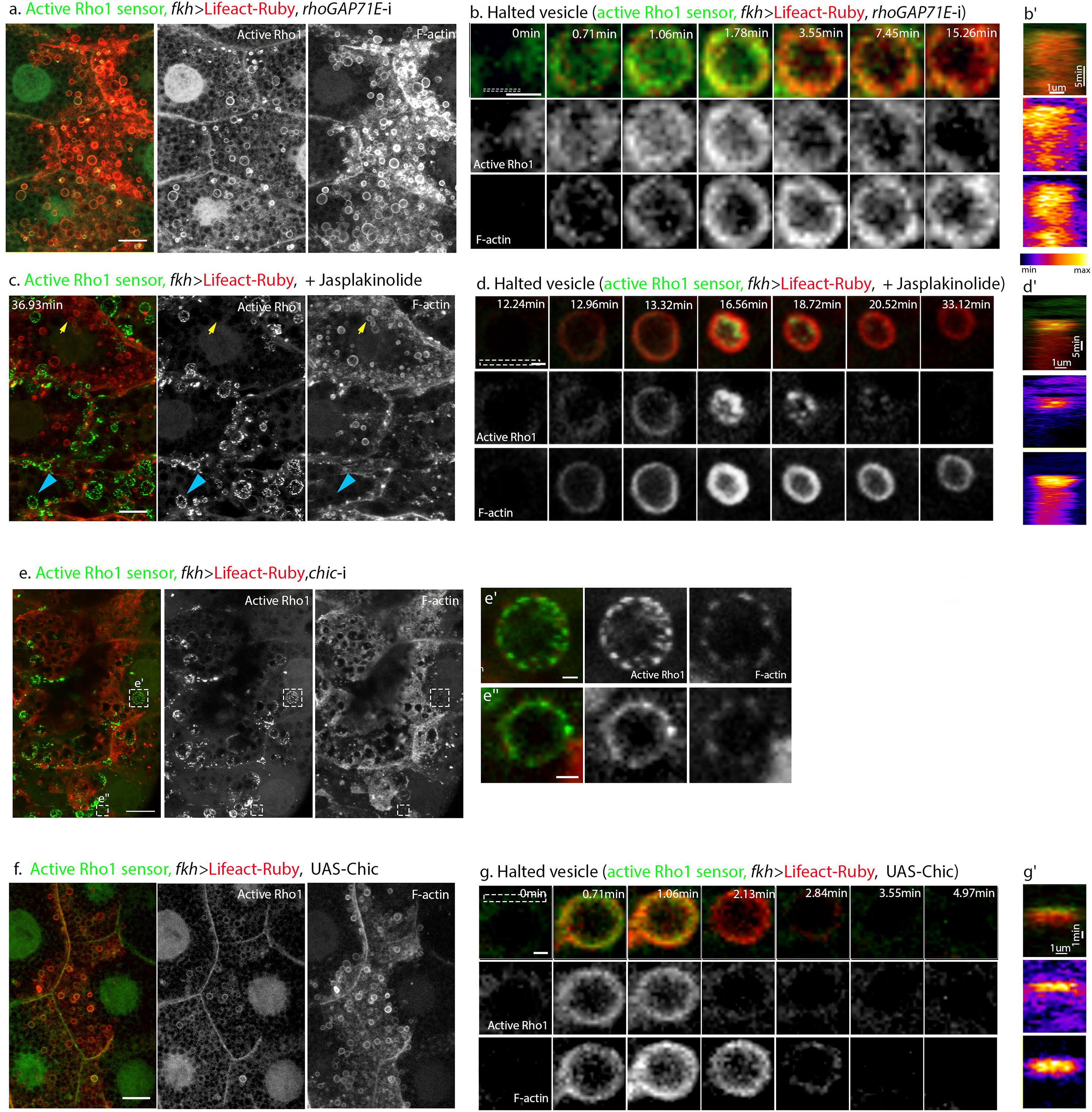
Actin disassembly is essential for vesicle contraction. **(a)** Salivary glands expressing the active Rho1 sensor (green) and Lifeact-Ruby (red) along with *rhoGAP71E-*i display accumulation of halted vesicles and sustained active Rho1 and F-actin, suggesting that RhoGAP71E-dependent Rho1 inactivation is necessary for actin coat disassembly and for proper vesicle contraction (see also Movie S3). **(b)** Time course of sustained active Rho1 sensor (green) and Lifeact-Ruby (red) on a single halted vesicle from such a gland. **(b’)** Kymograph monitoring active Rho1 and F-actin dynamics at the vesicle surface (dashed-box area in initial panel of **(b)**). **(c)** Glands expressing the active Rho1 sensor (green) and Lifeact-Ruby (red), treated with the actin-stabilizing drug Jasplakinolide was added. Two types of contraction-halted vesicles can be observed in these glands. Initial fusion events generate halted vesicles that display a sustained actin coat, but lose active Rho1 (yellow arrows, see also **(d)**). Vesicles that fuse following prolonged exposure to the drug display sustained active Rho1 but fail to form an actin coat (blue arrows; see also Movie S4). **(d)** Time course of transient active Rho1 sensor (green) and sustained Lifeact-Ruby (red) on a single halted vesicle from the initial pool generated following drug treatment. Times indicated in (**c,d**) correspond to time elapsed since drug addition. (**d’)** Kymograph monitoring active Rho1 and F-actin dynamics at the vesicle surface (dashed-box area in initial panel of **(d)**). **(e-e”)** Salivary glands expressing the active Rho1 sensor (green) and Lifeact-Ruby (red) along with *chic-*i display fused contraction-halted vesicles that exhibit enriched active Rho1 but only a weak, punctate actin coat, suggesting that robust actin coat polymerization depends on Chic (see also movie S5). **(e’)** and **(e”)** show enlargements of boxed areas in **(e)**. **(f)** UAS-driven overexpression of Chic in glands expressing the active Rho1 sensor (green) and Lifeact-Ruby (red) leads to accumulation of contraction-halted vesicles (see also movie S6). **(g)** Normal time course of the active Rho1 sensor (green) and Lifeact-Ruby (red) on a single halted vesicle from such a gland. **(g’)** Kymograph monitoring active Rho1 and F-actin dynamics at the vesicle surface (dashed-box area in initial panel of **(g)**). Scale bars: 20μm in a, c, e, f. 2μmin b, d, e’, e”, g.

An alternative method to prevent actin turnover is by treatment with the microfilament-stabilizing drug Jasplakinolide (Bubb et al., 1994). Contraction-halted vesicles with a persistent actin coat were observed shortly after treatment, again demonstrating the importance of local actin turnover for actomyosin contraction (Figure 4c, d). Active Rho1 was not sustained on these vesicles, suggesting that additional input(s) besides persistence of the actin coat contribute to inactivation of Rho1. Notably, prolonged exposure to Jasplakinolide gave rise to a second class of halted vesicles, lacking an actin coat, likely caused by depletion of the pool of free actin monomers in the absence of actin turnover (Figure 4c, Movie S4). These vesicles displayed sustained active Rho1, similarly to what was observed upon addition of Lat-A and consistent with a role for F-actin in Rho1 inactivation (see Figure 2g, h).

In order to further examine the role of actin turnover in vesicle constriction, we interfered with actin polymerization dynamics by manipulating the levels of Chickadee (Chic), the *Drosophila* Profilin homolog (Cooley et al., 1992). Profilin is a key actin monomer-binding protein, which affects actin polymerization in a variety of ways, in different cellular contexts (Rotty et al., 2015; Shields et al., 2014; Suarez et al., 2015). Knockdown of *chic* led to formation of abnormally thin, punctate actin coats and an arrest of vesicle contraction (Figure 4e-e”, Movie S5), consistent with a role for Profilin-bound actin in mediating the Dia-dependent polymerization underlying coat assembly. Interestingly, overexpression of Chic, presumably increasing Dia-dependent polymerization, also caused defective squeezing, although an effect on coat structure was not immediately apparent (Figure 4f, Movie S6). These observations suggest that tight regulation of actin polymerization is necessary for proper actomyosin contraction.

### Branched actin is involved in regulation of actin disassembly

Previous work has identified the branched actin polymerization machinery, centered on the Arp2/3 complex, as an important mediator of glue vesicle contraction, although it is not essential for formation of the bulk of the actin coat (Rousso et al., 2016; Tran et al., 2015). Indeed, addition of the Arp2/3 inhibitor CK-666(Nolen et al., 2009) to secreting salivary glands led to accumulation of contraction-halted vesicles (Figure 5a, Movie S7). We wanted to test if branched actin also plays a role in the dynamic regulation of Rho1 activity. CK-666 was added to our experimental regime of Y-27632-treated glands expressing the active Rho1 sensor and Lifeact-Ruby. The halted vesicles in these glands showed both sustained active Rho1 and a sustained actin coat (Figure 5b-c, Figure S2c). This phenotype is similar to that observed following knockdown of *RhoGAP71E* (see Figure S2), suggesting that branched actin is also required for Rho1 inactivation.

**Figure 5:**
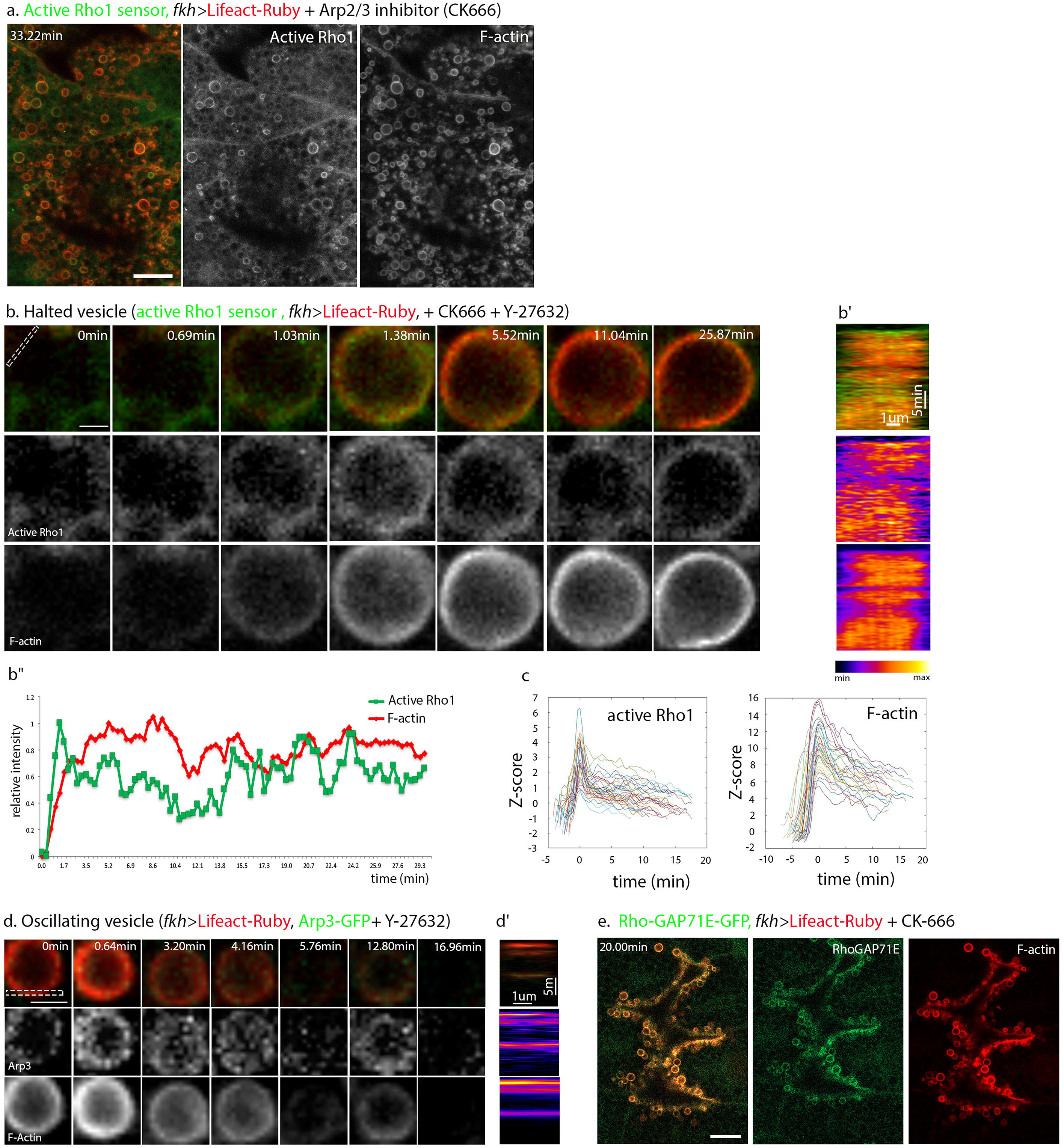
Branched actin is involved in regulation of actin disassembly. **(a)** Salivary glands expressing the active Rho1 sensor (green) and Lifeact-Ruby (red), and treated with the Arp2/3 complex inhibitor CK-666, which blocks branched actin polymerization, show accumulation of contraction-halted vesicles enriched for active Rho1 and F-actin (see also Movie S7). Time indicated corresponds to time elapsed since drug addition, **(b)** Sustained active Rho1 and a sustained F-actin coat are displayed by a single contraction-halted vesicle from a gland expressing the active Rho1 sensor (green) and Lifeact-Ruby (red), and treated with both Y-27632 and CK-666, **(b’)** Kymograph monitoring active Rho1 and F-actin dynamics at the vesicle surface (dashed-box area in initial panel of **(b)**). **(b’’)** Graph plotting relative intensity of active Rho1 (green) and F-actin in the area monitored in the kymograph **(b’)** over time. Intensity is normalized to the highest value within each channel. Note that both active Rho1 and actin rise sharply and are sustained on the vesicle for over 20 minutes, similar to what was observed following knockdown of *RhoGAP71E*. **(c)** Dynamic profiles over time of active Rho1 and F-actin in 36 vesicles from a single contraction-halted gland with added CK-666. Z-score is defined as the number of standard deviations that the signal intensity differs from background levels. Note that active Rho1 and actin rise sharply and show an initial partial decline, but are sustained above basal levels over many minutes (compare timescale to Figure 1f). **(d)** Contraction-halted glands expressing Arp3-GFP (green) and Lifeact-Ruby (red) display coordinated oscillations of Arp3-GFP and the actin coat, similarly to active Rho1 and RhoGAP71E-GFP, suggesting that the Arp2/3 machinery is part of this shared regulatory circuitry. **(d’)** Kymograph monitoring Arp3 and F-actin dynamics at the vesicle surface (dashed-box area in initial panel of **(d)**). **(e)** Glands expressing RhoGAP71E-GFP (green) and Lifeact-Ruby (red) and treated with CK-666 show accumulation of contraction-halted actin-coated vesicles. RhoGAP71E is also recruited and sustained on fused vesicles. Although Arp2/3 activity appears to be important for the activity of RhoGAP71E (since CK-666 treatment results in sustained active Rho1), RhoGAP71E-GFP recruitment to the vesicle is not dependent on branched actin polymerization. Scale bars: 20μm in a, e. 2μm in b, d.

Elements of the Arp2/3 complex, particularly Arp3, are enriched on fused vesicles (Tran et al., 2015). To assess Arp2/3 dynamics relative to the actin cycle, we expressed Arp3-GFP in salivary glands and followed its recruitment to contraction-halted vesicles. Indeed, Arp3-GFP enrichment on fused, contraction-halted vesicles oscillates in correlation with F-actin (Figure 5d), similarly to the active Rho1 sensor and RhoGAP71E-GFP (see Figure 3), consistent with the notion that branched actin polymerization is also part of this shared circuitry.

One possibility suggested by these observations is that branched actin inhibits Rho1 by facilitating the recruitment of RhoGAP71E to the fused vesicles. However, appearance of RhoGAP71E-GFP on vesicles was not affected by addition of the Arp2/3 inhibitor, indicating that recruitment of RhoGAP is not dependent on branched actin assembly (Figure 5e). These results suggest therefore that branched actin polymerization may influence the process by affecting RhoGAP71E activity rather than its recruitment.

## Discussion

Exocytosis of adhesive glycoproteins into the lumen of *Drosophila* larval salivary glands serves as an established model for the mechanistic basis of vesicle secretion in tubular organs(Tran and Ten Hagen, 2017). Once the large, cargo-filled secretory vesicles fuse with the apical membrane of the gland epithelial cells, the coordinated recruitment of the actin nucleation machinery and Myosin II is essential for vesicle contraction and subsequent content expulsion into the gland lumen (Rousso et al., 2016; Tran et al., 2015). In this work, we followed the dynamics of actin coating of individual salivary gland secretory vesicles, and identified a cycle of actin coat assembly and disassembly that is independent of myosin and vesicle constriction. We determined that the time course of a single actin coat assembly/disassembly cycle is roughly similar to that of a contracting vesicle, allowing it to operate within the normal timeframe of vesicle contraction. The molecular events underlying the actin cycle are set in motion upon fusion of the vesicle to the apical membrane and recruitment of activated Rho1 to the vesicle surface, triggering both the actin-polymerizing activity of the formin Dia and the recruitment of Myosin II (Rousso et al., 2016). The assembled F-actin coat on the fused vesicle then recruits RhoGAP71E, leading to the inactivation of Rho1, resulting in actin coat disassembly. The branched actin nucleator Arp2/3 is also recruited to the fused vesicle, and is essential for the shutdown of Rho1. Figure 6 summarizes the actomyosin cycle that takes place on the fused vesicle.

**Figure 6:**
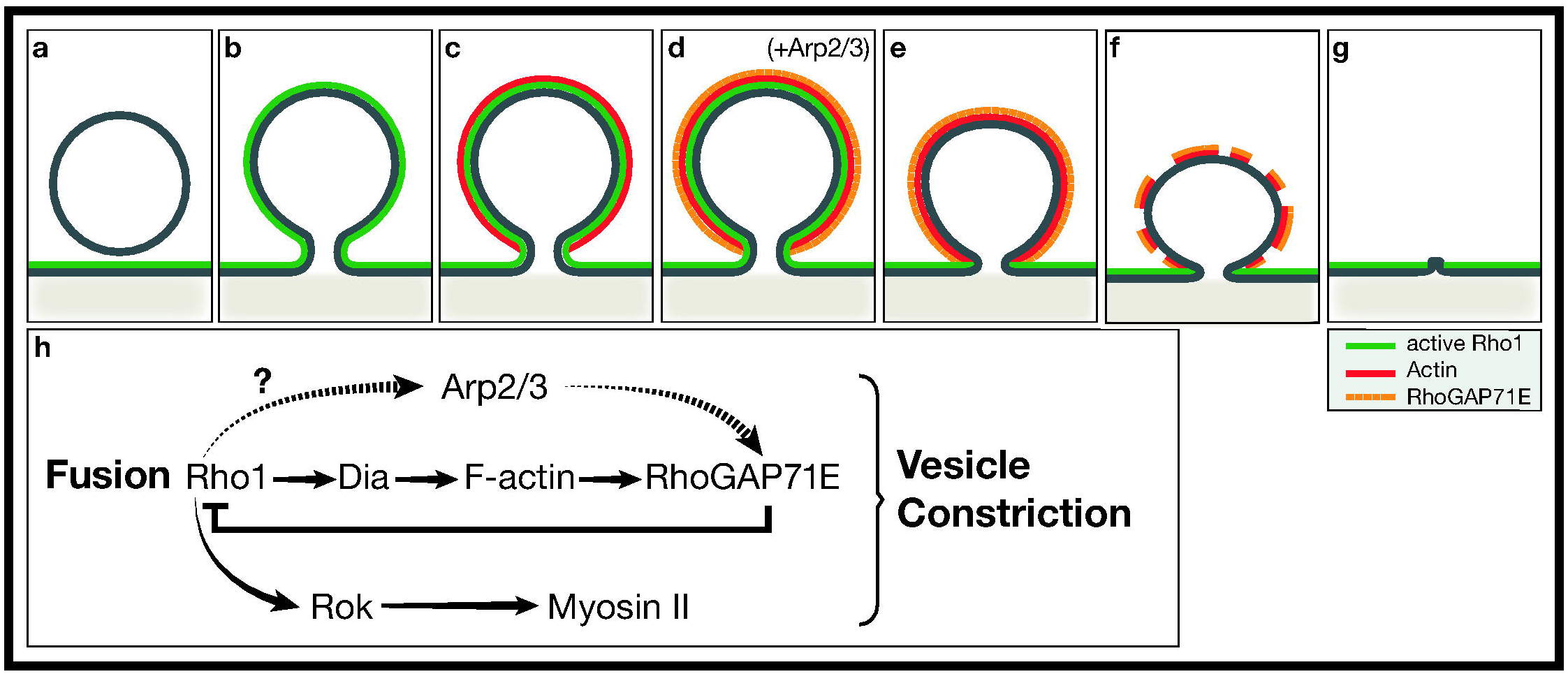
Coordinated formation and disassembly of a contractile actomyosin network mediates content release from large secretory vesicles. A cycle of actin assembly and subsequent depolymerization occurs on a contracting secretory vesicle. The composition of the secretory vesicle membrane is different from the apical salivary gland membrane **(a)**. Following vesicle fusion to the apical membrane, active Rho1 is enriched on the vesicle surface **(b)**. Active Rho1 then triggers assembly of linear actin via Dia **(c)**, which in turn recruits RhoGAP71E **(d)**. RhoGAP71E inactivates Rho1 **(e),** resulting in actin disassembly **(f)**, and allowing for actomyosin-mediated vesicle constriction **(g)**. Arp2/3-mediated branched actin nucleation is also recruited to the fused vesicle **(d)**, and is essential for Rho1 inactivation. **(h)** Fusion of the vesicle recruits activated Rho1 from the apical membrane, leading in parallel to the actin nucleation and depolymerization cycle described above, to the recruitment of Arp2/3, and to the recruitment of Rok which triggers Myosin II activation. While Rho1 regulates the recruitment of Myosin II to the vesicle, the dynamics of the actin coat are myosin-independent. Activities of all three arms are essential for vesicle constriction.

Importantly, we found that disassembly of the actin coat is a critical aspect of the secretory process, implying that local actin turnover is essential for actomyosin-mediated vesicle contraction. Thus, upon RhoGAP71E knockdown, persistent Rho1 activity caused stabilization of the actin coat, leading to failure of vesicle constriction. Similarly, chemical stabilization of the actin coat or enhancement of actin polymerization by overexpression of Profilin also abolished the capacity of the vesicles to squeeze. Actin turnover has been shown to be essential for actomyosin contraction in diverse cellular processes, including cytokinesis in *C. elegans* and fission yeast (Carvalho et al., 2009; Vavylonis et al., 2008), and adjustable adhesion between epithelial cells in *Drosophila* (Jodoin et al., 2015). The requirement for actin disassembly in constriction of secretory vesicles described in this work emphasizes the importance of regulation of actin dynamics across diverse systems.

Why is actin turnover essential for glue vesicle constriction? The actomyosin network in this system faces a significant challenge, in that it must contract and squeeze a large vesicle with a viscous content within a short time frame. We can envision two ways where actin turnover may play a significant role in facilitating this process. First, actin turnover may be important for dynamic reorganization of myosin as a vesicle contracts. Recent theoretical work has suggested that actin turnover may be important to randomize actomyosin arrays, in order to avoid stabilization of configurations that are inefficient for contraction (Rubinstein and Mogilner, 2017). For each myosin motor, movement along a single linear actin track may be insufficient for contraction of a large surface area such as that of the vesicle. Dissociation of the actin network may thus allow a motor that has already moved along a given actin track to reorganize along a new track and carry out a new cycle of movement.

Another possible explanation relates to the dynamics of actin nucleation that dictate the balance between linear and branched actin polymerization, since both are essential for vesicle contraction (Rousso et al., 2016; Tran et al., 2015). Under circumstances where monomeric actin (G-actin) is a limiting resource, actin bound by Profilin favors formin-based linear actin over Arp2/3-mediated branched actin assembly (Burke et al., 2014; Suarez et al., 2015). The availability of free G-actin monomers near the fused vesicles may be limiting in this system, as demonstrated by the inability of newly fused vesicles to form an actin coat, once F-actin was chemically stabilized (Fig. 4). Thus, following fusion, the nucleation of actin may be carried out primarily by Dia. Once Rho1 activation is blocked by RhoGAP71E, the newly-available actin monomers can be channeled to the branched actin nucleation machinery. This model suggests a self-regulatory cascade, where the recruitment of RhoGAP71E leads to activation of the branched actin-nucleation machinery. While the specific role of actin turnover in this system remains an open question, this work demonstrates the need for local actin turnover for proper actomyosin function, and suggests a mechanism for its self-organized regulation.

The feedback cycle of actin assembly and disassembly on the secretory vesicles, mediated by Rho1, Dia, and RhoGAP71E, could represent a broader paradigm in actomyosin dynamics. The capacity of the product, F-actin, to trigger negative feedback on its own synthesis via recruitment of a RhoGAP, may be used in diverse biological scenarios and be adjusted to the particular requirements of each setting. Several examples for actin negative feedback circuits were recently reported in early embryos. *Bement* et al. (2015) (Bement et al., 2015) monitored the surface of starfish and frog oocytes, and identified multiple successive waves of Rho activation and actin nucleation, followed by inactivation and disassembly, respectively. They speculated that these waves keep the oocyte surface undetermined, until a prominent signal for the localized recruitment of the actomyosin machinery is provided after fertilization, to generate the first cytokinetic ring. *Robin* et al. (2016) (Robin et al., 2016) followed spikes of actin nucleation and disassembly during pulsed contractility of the early *C. elegans* embryo, and identified two RhoGAP proteins that are recruited to the sites of actin nucleation and keep the cell surface plastic. An analogous circuitry was recently shown to regulate cell contractility in cell culture (Graessl et al., 2017). In this case, the actin nucleation and disassembly cascade was coupled to the activity of Myosin II, which may allow spontaneous patterns of subcellular contractility to explore local mechanical cues. A similar actin assembly and disassembly feedback module may contribute to other processes where dynamic waves of actin have been observed, such as in cell motility (Barnhart et al., 2017; Xiong et al., 2010), or formation of axonal protrusions (Katsuno et al., 2015).

Secretion of viscous materials from a variety of epithelial glands requires exertion of active forces, to facilitate content release from large secretory vesicles after their fusion with the apical cell membrane (Geron et al., 2013; Miklavc et al., 2012; Nightingale et al., 2011). Formation of an actin coat around each vesicle and recruitment of myosin is a common feature of these systems. Future work should examine whether the dynamic behavior of the actin coat and the functional significance of its disassembly, characterized here in the context of the *Drosophila* larval salivary gland, is also operating in other secretory systems.

## Materials and Methods

### *Drosophila* genetics

The following *Drosophila* lines were used (“B” designates BDSC stock number): UAS–LifeAct–Ruby (B-35545) (Hatan et al., 2011), Sgs3–GFP (Biyasheva et al., 2001) (B-5884), Sqh–GFP (Royou et al., 2004) (B-57145), *Ubi-AniRBD::GFP* (Rho1 sensor (Munjal et al., 2015)), UAS-Chickadee (from L. Cooley, Yale University), UAS-Arp3-GFP (Hudson and Cooley, 2002) (B-39721), UAS-*chic*-RNAi^HMS00550^ (B-34523). RNAi lines used for the RhoGAP screen are described in Table S1. *fkh-*Gal4 was used to drive expression in salivary glands (Maybeck and Roper, 2009). For the RhoGAP screen, flies were grown at 25°C. For glands expressing RNAi lines displayed in figures and used for further analysis, crosses were shifted to 29°C at 2^nd^ instar larval stage, overnight, for maximum effect of RNAi. Driver controls grown under the same regimen showed normal secretion.

RhoGAP71E-GFP was generated by injection of the plasmid: *Splice phase 0 EGFP-FIAsH-StrepII-TEV-3xFlag* plasmid, into the fly line Mi{MIC}RhoGAP71E^MI09170^, as described in (Venken et al., 2011).

### Culturing third-instar salivary glands

Following dissection in Schneider’s medium, several salivary glands were placed in a chamber slide (1μ-Slide 8 well Microscopy Chamber, Ibidi) containing 200□μl of medium. Ecdysone (5□μM 20-hydroxyecdysone, Sigma) was added to the medium immediately. For imaging, glands were trasnferred to a 35mm #0 Glass Bottom Dish, with 14mm Bottom Well (Cellvis D35-14-0-N), containing 200□μl fresh medium. Glands were submerged and arranged on the bottom of dish using forceps. The chamber slide was placed at room temperature, with light shaking for 2-3□h. Secreting glands were identified by their expanded lumen as viewed under a stereomicroscope before imaging. In many cases, salivary glands were dissected and imaged immediately during their natural glue-secretion phase.

### Time-lapse imaging and post-imaging processing

Data were acquired with an LSM800 confocal microscope system (Zeiss), using ×40/1.2 (water immersion) or ×20/0.8 (air) objectives and ×1 digital zoom. Typically, 18–20 *z*-slices were imaged at intervals of 0.5□μm (×40) with a temporal resolution of 20-40 seconds per frame. Data acquisition for the high time resolution data of RhoGAP71E-GFP (Figure S3) was carried out on a single z-slice, with a temporal resolution of 4.5 seconds per frame. Kymographs were created using the “reslice” function in Fiji. Post-imaging processing was carried out using Fiji and Adobe Photoshop CS3 for cropping and adjustment of Brightness/Contrast, for visualization purposes only.

### Drug treatment and dye injection

Each gland was imaged for at least 3 minutes before drug treatment to ensure overall gland health and commencement of secretion. With image acquisition stopped, ROCK inhibitor (Y-27632, 50μM, Sigma), Arp2/3 inhibitor (CK-666, 0.5mM, Sigma), Latrunculin-A (Lat-A, 1μM), or Jasplakinolide (Jasplak, 2 μM) were added to medium with a micropipette, directed near but not touching the sample, and mixed slightly in medium with pipette tip, after which acquisition recommenced. For dye injection experiments, Dextran–TMR (10,000□MW, 1□mg□ml^−1^ in Schneider’s medium, Molecular Probes) was injected into salivary glands during the glue-secretion phase using a Femtojet express microinjector and custom-made capillaries. Immediately following injection, glands were taken for imaging as described above.

### Characterization of signal dynamics

For individual representative vesicles, intensity measurements were made by plotting the mean intensity values of a line spanning the vesicle kymograph, for each channel, through time. Each channel was normalized to its own highest (1) and lowest (0) point, in order to compare temporal dynamics between active Rho1 and actin. Plots were created in Excel.

Characterization of pooled data from many vesicles was accomplished as follows:

**Manual annotation of vesicles:** Using Fiji, vesicles were manually identified in the field of view based on the actin channel. A time lapse sequence was cropped, starting from at least 2 time frames before actin coat appearance, and continuing for the entire duration that the vesicle could be tracked. To counter movement of vesicles in the focal plane (z-position), the z-slice where vesicle’s intensity was brightest was manually set for each time point. A vesicle’s bounding box was manually drawn based on the dimensions of the vesicle with the *frame of reference*, manually defined as the first appearance of signal in the actin channel. Cropped time-lapse images of vesicles were rotated as needed, to ensure that the bottom portion of the vesicle remained unobscured by neighboring vesicles. **Vesicle alignment:** In order to counter vesicle movement in the XY plane, each time frame within a vesicle’s time-sequence was aligned with respect to the frame of reference, using Matlab. A translation (Δx, Δy) was calculated between each frame in the sequence and the frame of reference as follows. Morphological dilation of 3 pixels expanded the annotated bounding box so as to include background pixels for the registration. A Gaussian filter with a width of 5 pixels and σ = 0.7 was applied to smooth both images and reduce high-frequency artifacts. The optimal translation maximized the Bhattacharyya coefficient of the normalized images with a maximal translation of 10 pixels (3.33 µm) in each direction. **Quantifying a vesicle’s dynamics:** A *reference line* was calculated for each time-frame and was used to quantify a vesicle’s dynamics over time. The reference line was defined as the horizontal 1-pixel wide mask that maximizes the actin channel’s median intensity within the bottom 5 rows from the vesicle’s bounding box after being aligned according to the reference frame. All vesicles were manually examined for mistakes in reference-line vesicle identification, such as labeling of neighboring vesicles. For each channel, the mean intensity of the five brightest pixels in the reference line at each frame was recorded as the *vesicle signal*. The vesicle’s signal was smoothed in time with a Gaussian filter of window size 5 pixels and σ = 2.5. To standardize vesicle intensities and time dynamics within and across experiments we calculated a *background intensities model* for each experiment. The model was designed to capture the image background pixel intensities statistics and use it for (1) correcting for photo-bleaching effects at the single experiment level; and (2) normalize vesicle intensities so that vesicles from different experiments could be pooled and compared. The signal of the first two time points for every vesicle were recorded as ‘background’. Background signal of all vesicles were pooled. In the cases were photo-bleaching was identified, i.e., a significant negative Pearson’s correlation between the background signal and time, we fitted a linear model that was used to correct the vesicle signal. Otherwise, the raw vesicle signal was used. The mean (denoted μ) and standard deviation (denoted σ) of the background was used to normalize each vesicle’s signal *x* according to the z-score measure: *x*^*norm*^ = (*x* − μ)/σ, i.e., the variation from the mean background in units of standard deviations that can be pooled and compared across experiments. **Quantifying oscillations:** Peaks were detected in the normalized vesicle signal such that peak intensity value is at least 3 standard deviations above the background and there are at least 140 seconds (7 frames) between two peaks. We filtered out peaks where the drop-off values on both sides before encountering a larger value are less than 1.5 standard deviation. We then scored the number of oscillations (e.g. initiations of actin cycles) in each vesicle, for the Y-27632-treated glands. In order to correct for bias of the limited durations that vesicles could be tracked, we scored the number of oscillations only in vesicles that were tracked for at least 15.5 minutes, defined by the 90^th^ percentile of times between oscillation peaks (Figure 3b), plus 3 seconds. **Quantifying time course of signal:** A single actin/active Rho1 cycle was calculated for each vesicle, at each channel, as follows. Vesicle background was defined as the mean z-score signal of the first 2 time points. The onset of the cycle was defined as the first time point where the signal intensity exceeds 2 standard deviations (σ) above the vesicle background. Note, that σ is calculated using the background from the entire gland, as described above. The cycle ends once the signal falls below 2 σ in relation to the vesicle’s background. A vesicle is reported to be “sustained” if the signal never returns below this threshold. Cycle time for sustained vesicles is defined as the duration of the video. This characterization was used for calculation of the distribution of time course of a single actin/active Rho1 cycle (see Figure S2). For calculation of the time lag of RhoGAP71E-GFP relative to actin, we defined the onset of each signal within a vesicle as described above, and subtracted time of actin signal onset from time of RhoGAP71E signal onset (see Figure S3c).

BoxplotR was used to generate box plots (Spitzer et al., 2014). Non-parametric statistical tests were carried out using Matlab where specified. Quantification of the number of vesicles showing sustained active Rho1 for at least 10 minutes on Lat-A treated glands was performed by manual scoring of vesicles on maximum intensity projections of images.

### Reproducibility

At least 3 salivary glands (from 3 different larvae) were examined for each genetic background and/or drug treatment, and representative images/videos are shown. Quantifications of specific vesicle phenotypes were made on representative videos. In addition, the dynamics of active Rho1 in the Lat-A experiments varied between glands, with 3 out of 5 glands exhibiting vesicles with sustained active Rho1.

## Acknowledgments

We thank Dana Meyen and Agur Wiskott for creating reagents and help with experiments. We thank L. Cooley and T. Lecuit for the UAS-Chickadee and active Rho1 reporter strain, respectively, and the BDSC and VDRC stock centers for *Drosophila* lines. We are grateful to members of the B-Z.S. laboratory for stimulating discussions. This work was supported by an Israel Science Foundation grant to E.D.S. and B-Z.S, and a National Insitute of Health Grant to Gaudenz Danuser supporting A.Z. (P01 GM103723). B-Z.S. is an incumbent of the Hilda and Cecil Lewis chair in Molecular Genetics.

## Author contributions

D. Segal conceived the research concept, developed methodologies, carried out and analyzed experiments, and wrote the manuscript. A. Zaritsky designed and carried out computational analysis of data. E.D. Schejter and B. Shilo conceived the research concept, supervised experiments, and wrote the manuscript.

